# Prolonged Withdrawal from Escalated Oxycodone is Associated with Increased Expression of Glutamate Receptors in the Rat Hippocampus

**DOI:** 10.1101/2020.12.20.423680

**Authors:** Aaron J. Salisbury, Christopher A. Blackwood, Jean Lud Cadet

**Affiliations:** National Institute on Drug Abuse, Molecular Neuropsychiatry Branch, National Institutes of Health, Baltimore, MD, USA

**Keywords:** AMPAR, NMDAR, GRM, OUD, Hippocampus

## Abstract

People suffering from opioid use disorder (OUD) exhibit cognitive dysfunctions. Here, we investigated potential changes in the expression of glutamate receptors in rat hippocampi at 2 hours and 31 days after the last session of oxycodone self-administration (SA). RNA extracted from the hippocampus was used in quantitative polymerase chain reaction (qPCR) analyses. Rats, given long-access (9 hours per day) to oxycodone (LgA), took significantly more drug than rats exposed to short-access (3 hours per day) (ShA). In addition, LgA rats could be further divided into higher oxycodone taking (LgA-H) or lower oxycodone taking (LgA-L) groups, based on a cut-off of 50 infusions per day. LgA rats, but not ShA, rats exhibited incubation of oxycodone craving. In addition, LgA rats showed increased mRNA expression of *GluA1-3 and GluN2a-c* subunits as well as *Grm3*, *Grm5*, *Grm6* and *Grm8* subtypes of glutamate receptors after 31 days but not after 2 hours of stopping the SA experiment. Changes in *GluA1-3, Grm6, and Grm8* mRNA levels also correlated with increased lever pressing (incubation) after long periods of withdrawal from oxycodone. More studies are needed to elucidate the molecular mechanisms involved in altering the expression of these receptors during withdrawal from oxycodone and/or incubation of drug seeking.

## 1 Introduction

The opioid epidemic that includes the abuse of oxycodone is associated with large numbers of overdose-related deaths [Wilson et al., 2020]. Oxycodone is a semisynthetic opioid analgesic prescribed to patients suffering from moderate to severe pain [Riley et al., 2008]. Oxycodone use disorder (OUD) is a chronic relapsing disorder characterized by compulsive drug taking despite adverse life consequences [DSM-V, 2013]. In people with OUD, neurocircuits in the brain’s reward systems that control hippocampus-mediated cognitive processes including learning and memory [Cadet et al., 2014] are altered [Koob and Volkow, N. D. 2010]. Cognitive processes are indeed affected in patients who abuse opioids [Allegri et al., 2019; Kroll et al., 2018].

Although the hippocampus is essential for cognitive functions that can be disturbed in substance use disorders (SUDs), it has received much less attention than other brain regions such as the nucleus accumbens or dorsal striatum in studies involving animal models of SUDs. Nevertheless, the hippocampus has been shown to be important in the regulation of drug intake [Glick and Cox, 1978; Chambers and Taylor, 2004; Brady et. al, 2010] and to mediate context- and cue-induced reinstatement of drug taking after withdrawal [Fuchs et. al, 2005; Rogers et. al, 2007]. Importantly, alcohol and opioid exposure negatively impact adult hippocampal neurogenesis [Zhang et. al, 2016] and enhances long term potentiation (LTP) [Elahi-Mahani et. al, 2018]. Furthermore, there is evidence to show that the strength of hippocampal inputs into the nucleus accumbens can bidirectionally drive motivation for rewarding stimuli [Legates et. al, 2018]. While these studies have shown a significant role for the hippocampus in mediating drug taking and re-instatement, there is not enough research that documents the effects of opioid drugs on gene expression in the hippocampus. In order to develop more effective opioid addiction treatments, it is necessary to identify molecular neuroadaptations that occur in the hippocampus during long-term exposure and withdrawal from these drugs. To reach these aims, we have used a rat oxycodone self-administration model to probe the potential molecular changes that occur in that brain region.

The present study was designed to identify potential changes in the mRNA expression of several glutamate receptor subunits in the hippocampus of rats that had been exposed to oxycodone during drug self-administration experiments. So far, there had been no studies that examined changes in the expression and/or compositions of α-amino-3-hydroxy-5-methyl-4-isoxazolepropionic acid receptors (AMPARs) and N-methyl-D-aspartate receptors (NMDARs), both of which are important for the induction and maintenance of long-term potentiation (LTP), a process that is impacted by opioids [Portugal et al, 2014; Terman et al, 1994]. It is also to be noted that metabotropic glutamate receptors (mGluRs) have also been implicated in animal models of substance use disorders (SUDs) [Olive 2009].

Herein, we report that long-term withdrawal from long-access to oxycodone is associated with selective increases in AMPAR and NMDAR subunits glutamate receptors in the rat hippocampus. Some subunits of Group I and Group III metabotropic receptors were also affected.

## 2 Experimental Procedures

### 2.1 Subjects

Male Sprague–Dawley rats, (Charles River Laboratories, Raleigh, NC, USA) weighing 350– 400 g, were housed singly prior to surgery on a 12-hours (h) light/dark cycle and had food and water freely available. All procedures were performed according to guidelines outlined in the eighth Edition of National Institutes of Health (NIH) Guide for the Care and Use of Laboratory Animals and were approved by the local National Institute of Drug Abuse Intramural Research Program, Animal Care and Use Committee (ACUC).

### 2.2 Intravenous Surgery and Self-Administration Training

Animals were surgically implanted with intravenous jugular catheters (Cadet et al., 2017, Blackwood et al., 2019a,b). An intraperitoneal injection of buprenorphine (0.1 mg/kg) was given to each rat to manage pain following surgery and were allowed 1 week of recovery before beginning self-administration. Rats were trained in self-administration chambers located inside sound-attenuated cabinets and controlled by a Med Associates System (Med Associates, St Albans, VT). Rats were housed in these chambers for the duration of the experiment. Rats were randomly assigned to either saline (Sal) or oxycodone groups. Oxycodone-assigned rats were trained to self-administer oxycodone-HCL (NIDA Pharmacy, Baltimore, MD) using short-access and long-access paradigms (Fig 1). Short-access (ShA) rats were trained for one 3-h daily session for the course of the experiment. Long-access (LgA) rats were trained for a single 3-h daily session during the first week of self-administration, two 3-h daily sessions with a 30 minute break in between sessions during the second week, and three 3-h daily sessions with a 30 minute break between sessions for the remainder of the self-administration. Lever presses were reinforced using a fixed ratio-1 with a 20-s timeout accompanied by a 5-s compound tone-light cue. Rats self-administered oxycodone at a dose of 0.1 mg/kg per infusion over 3.5-s (0.1 ml per infusion. The lever was made available and cue was presented along with an oxycodone infusion to signal the start of the session. At the end of each 3-h session and at the end of the day, the tone-light cue was turned off and the levers retracted. Saline rats were assigned to either a ShA or LgA training paradigm as well and received a 0.1 ml of 0.9% saline per infusion. After the last day of training, some rats were euthanized 2-h after the last self-administration session whereas other rats were returned to the animal vivarium and individually housed with no access to oxycodone during which time they participate in drug seeking tests under extinction conditions. Briefly, rats underwent 3-hour cue-induced drug seeking tests on withdrawal days 5 (WD5) and 30 (WD30) during which time presentations of the cue and lever pressing were not accompanied by any oxycodone infusion.

**Figure 1:**
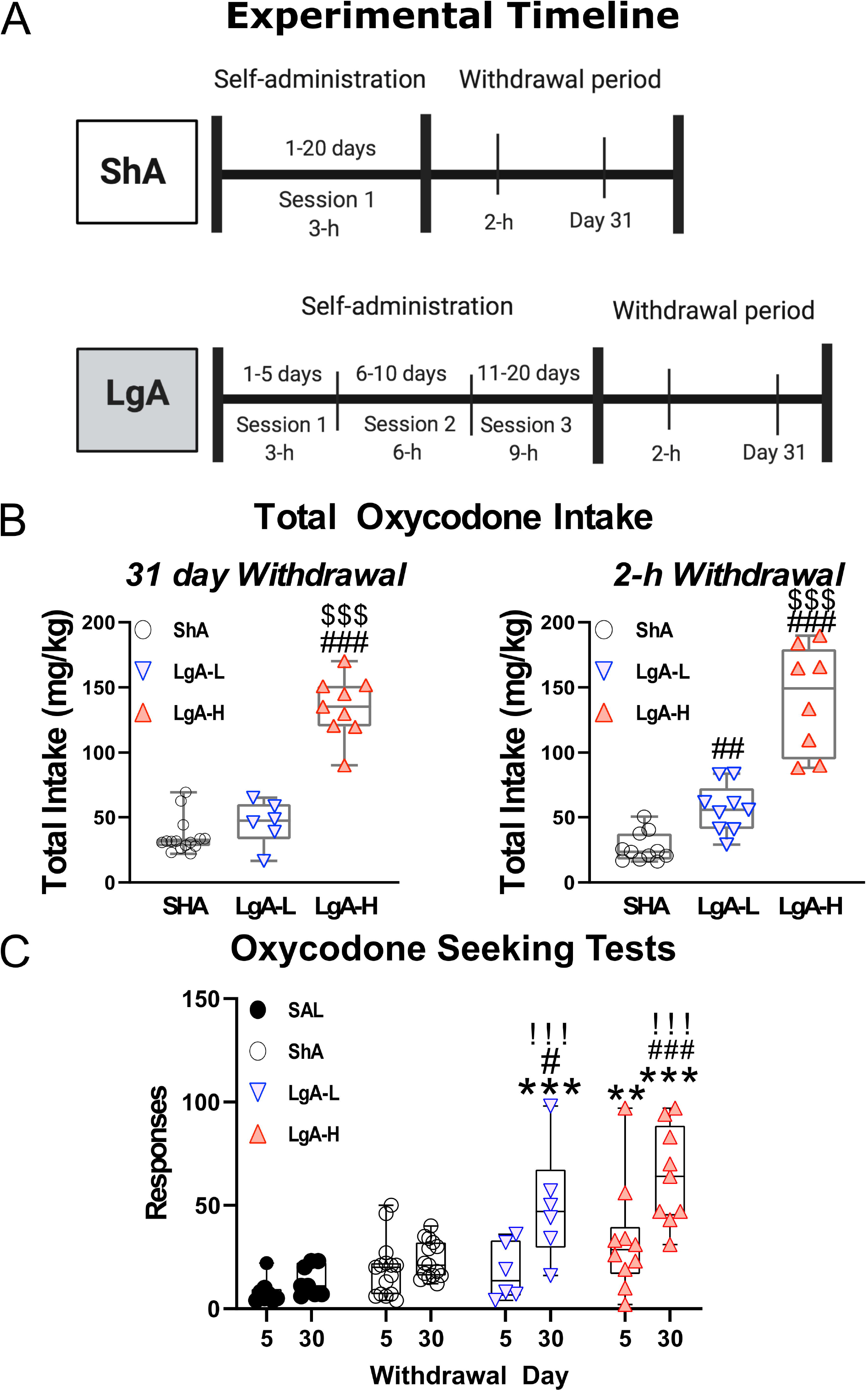
Experimental timeline and oxycodone intake by treatment group. **(A)** Experimental Timeline of oxycodone self-administration for ShA and LgA groups. **(B)** Shows the sum total oxycodone intake for each group from all three daily sessions over the course of the experiment **(C)** Shows the lever pressing during extinction tests on day 5 and 30 of withdrawal. *, **, *** = p < 0.05, 0.01, 0.001, respectively, in comparison to saline rats; #, ### = p < 0.05, 0.001, respectively, in comparison to ShA rats; !!! = p < 0.001 compared to the same treatment group on day 5.

Rats that experienced catheter failure or became sick and unable to continue in the experiment were removed from the study and excluded from further analysis.

### 2.3 mRNA Extraction and Quantitative RT-PCR

Rats were euthanized either 2-h after the last self-administration session or 24-h after the last oxycodone seeking test. We chose the 2-hr time points based on our previous experiments with methamphetamine (Cadet et al., 2014, 2017) and oxycodone (Blackwood et al., 2020), in which we were able to identify changes in mRNAs coding for immediate early genes, potassium channels, or stress-related peptides. We chose 24 hours after the drug seeking test because we assumed that most of the effects of only lever pressing would have disappeared after 24 hours. Therefore, we thought it like that we would be measuring mainly the effects of prolonged oxycodone withdrawal. We have published other studies with methamphetamine using a similar approach (Daiwile et al., 2019).

The hippocampus was then dissected and isolated using coordinates (A/P − 5 to −7 mm bregma, mediolateral ± to 6 mm, D/V − 2 to −8 mm) according to Paxinos and Watson, 1998. Collected hippocampi were then used for RNA extraction using RNeasy Mini Kit (Qiagen, Valencia, CA). RNA preparation and RTqPCR experiments were performed as previously described [Cadet et al., 2017]. *B2M,* a gene coding for the class I major histocompatibility complex protein β-2-microglobulin was used as reference gene, as it has been used previously in rats given opioids as well as methamphetamine [Blackwood et al. 2018, Zoubková et al 2019]. In addition, as a matter of principle, we always make sure that the expression of any reference gene is not altered under the conditions of our experiments before using it as a reference gene. The results are shown as fold changes calculated as the ratios of normalized gene expression data for oxycodone SA groups compared to the saline group. All quantitative data are presented as means ± SEM. Primer sequences are listed in Table 1.

**Table 1:**
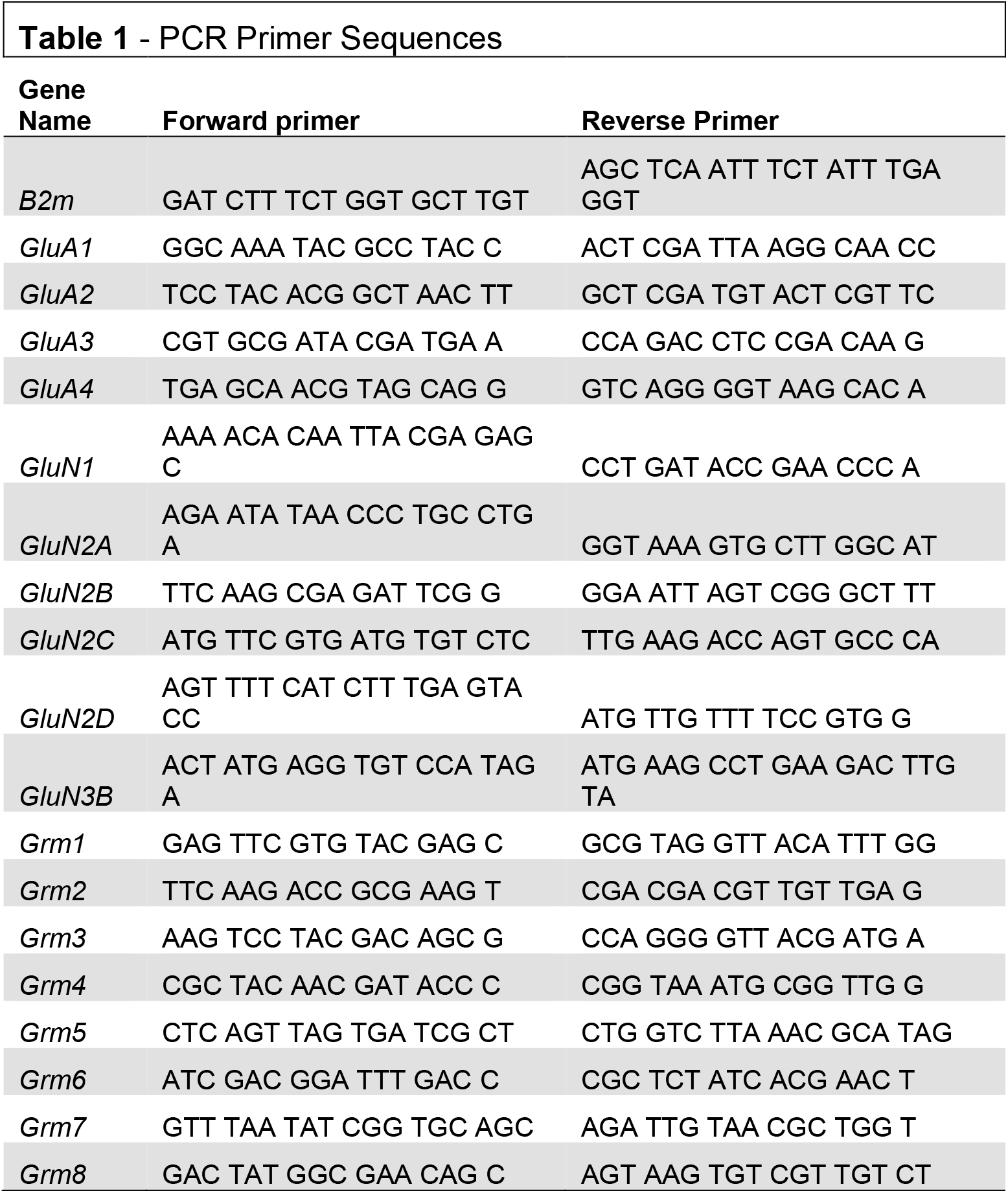
Shows RT-PCR primer sequences used in our experiments. *B2m* was used as a reference gene. Forward primers are complimentary to the anti-sense strand and reverse primers are complimentary to the sense strand.

### 2.4 Statistical Analyses

One-Way Analysis of Variance (One-Way ANOVA) was used to analyze the PCR data with the normalized fold change in mRNA levels as the dependent variable and the treatment group (SAL, ShA, LgA-L, LgA-H) as the independent variable. Outliers were excluded according to results of the Robust regression and Outlier removal Test (ROUT) method with Q = 1%. This was followed by Tukey’s post hoc test or Bonferroni post hoc test to look for significance between groups. Regressions were performed to look for correlations between oxycodone intake and mRNA expression. The null hypothesis was rejected at p < 0.05. all statistical tests were performed using GraphPad Prism version 8.4.2 (GraphPad Software, San Diego, CA).

## 3 Results

### 3.1 Rats Given Long-Access to Oxycodone Differentially Escalate Their Drug Intake

Fig. 1 shows the experimental timeline of the SA paradigm and the total amount of oxycodone taken by each rat over the course of the experiment. Rats were either given short-access to oxycodone or long-access (LgA) to oxycodone as described in the method section (Fig. 1A). LgA rats take significantly more oxycodone than the ShA rats. LgA groups could be divided into two further groups, long-access high (LgA-H) and long-access low (LgA-L), based on how they escalated their intake and how much oxycodone they ended up taking (Fig. 1B). Rats that took fewer than 50 infusions per day were put in the LgA-L group whereas the LgA-H consisted of rats that took more than 50 infusions per day. The LgA groups both showed incubation of craving during the drug seeking test on WD30 of withdrawal from oxycodone as reported previously (Blackwood et. al, 2018; 2019) (Fig. 1C).

### 3.2 Hippocampal AMPAR Subunit mRNAs are Differentially Regulated Following Withdrawal from Oxycodone SA

AMPA receptors participate in the regulation of neurotransmission, synaptic plasticity and LTP that are impacted by opioid exposure (Wang et al., 2004; Ortiz et al., 1995). Fig. 2 shows the effects of oxycodone intake and withdrawal on *GluA1-4*. mRNA levels. There were no significant differences in *GluA1* [F_(3, 30)_=1.641, p=0.2008], *GluA2* [F_(3, 29)_ = 0.479, p = 0.6994], or *GluA3* [F_(3, 30)_=1.013, p=0.4006] mRNA levels between groups at 2 hours after last oxycodone intake (Figs. 2A, D, and G). However, *GluA4* mRNA expression was significantly decreased [F (3, 29) =3.760, p=0.0214] (Fig. 2J) in the LgA groups compared to saline at that time. *GluA1* mRNA expression was significantly upregulated [F_(3, 30)_=3.730, p=0.0217] (Fig. 2B) in the LgA-H group compared to the ShA group and saline controls at 31 days. *GluA2* mRNA expression also showed significant increases [F_(3, 30)_=7.685, p=0.0006] (Fig. 2E) in both LgA groups compared to the ShA group and saline controls at 31 days. In addition, *GluA3* expression was significantly increased [F _(3, 30)_=5.000, p=0.0063] (Fig. 2H) in both LgA groups compared to the ShA group. In contrast, there were no significant changes in *GluA4* [F_(3, 30)_=2.108, p=0.1187] expression at that time (Fig. 2K). Interestingly, the changes in *GluA1*, *GluA2*, and *GluA3* mRNA levels were significantly positively correlated with increased lever pressing (incubation) after 31 days of withdrawal (Fig. 2C, F, I).

**Figure 2:**
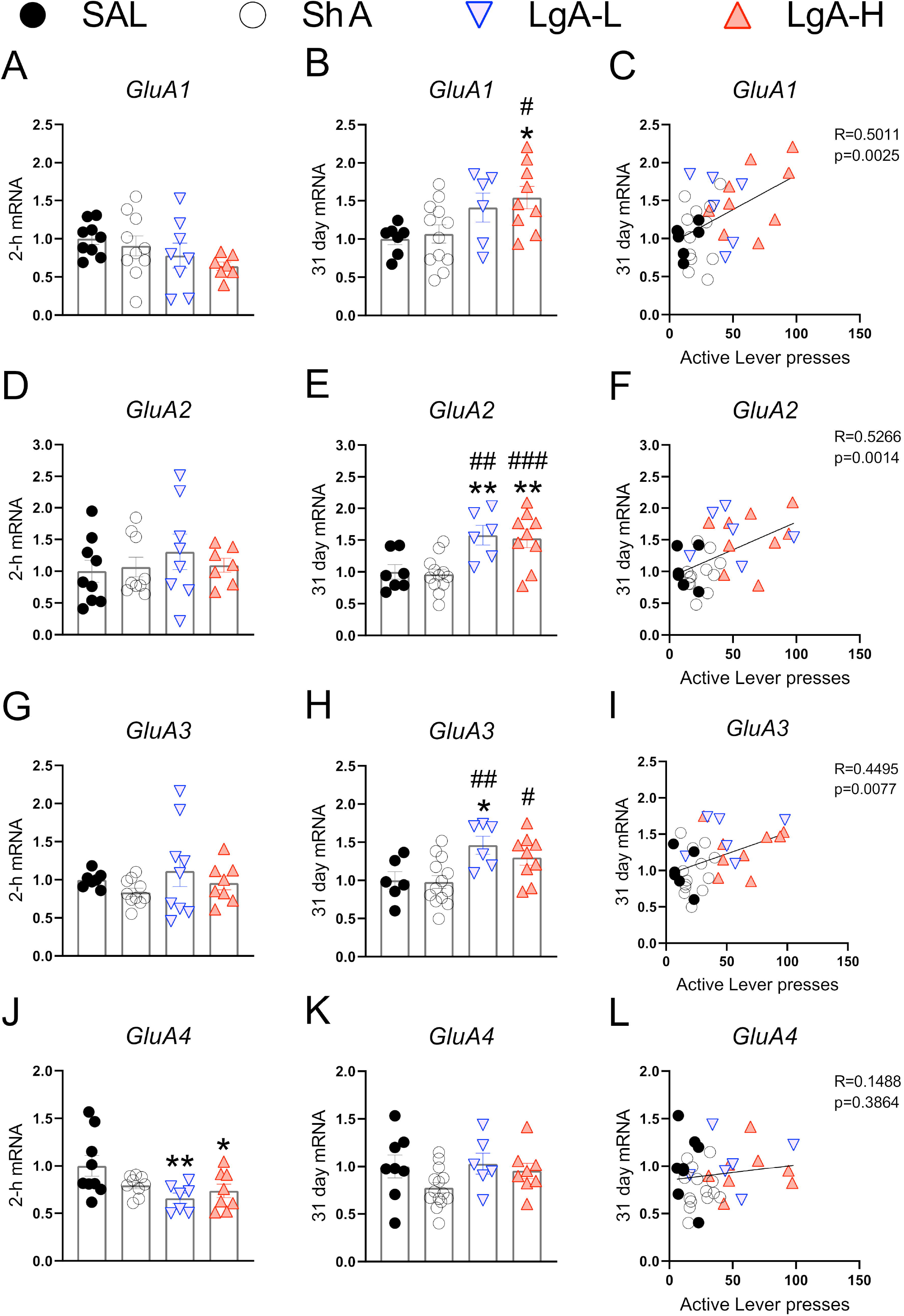
Rats undergoing long-term withdrawal from oxycodone showed increased expression of AMPA receptor mRNA. **(A-C)** *GluA1* mRNA expression is significantly increased in the LgA-H group compared to SAL at 31 days and showed a significant correlation with lever pressing on WD30. **(D-F)** *GluA2* mRNA expression is significantly increased in the LgA-L and LgA-H groups compared to the SAL and ShA groups at 31 days and showed a significant correlation with lever pressing on WD30. **(G-I)** *GluA3* mRNA expression is significantly increased in the LgA-L compared to the SAL and ShA grops as well as in the LgA-H group compared to the ShA group at 31 days and showed a significant correlation with lever pressing on WD30. **(J-L)** *GluA4* mRNA expression is significantly decreased in the LgA-L and LgA-H group compared to SAL at 2 hours and showed no significant correlation between 31 day mRNA expression and lever pressing on WD30. Key to statistics: *, **, = p < 0.05, 0.01, respectively, in comparison to Sal rats; #, ##, ### = p < 0.05, 0.01, 0.001, respectively, in comparison to ShA rats. Statistical Analyses were performed by One Way ANOVA followed by Fisher’s PLSD post hoc test, and correlation was tested by simple linear regression.

### 3.3 Selective Decreases in GluN Subunit Expression in LgA Rats after Drug Withdrawal

NMDA Receptors are also important regulators of synaptic plasticity and synaptic transmission [Lüscher & Malenka, 2012]. They work in tandem with AMPA receptors to facilitate synaptic transmission and regulate LTP [Lüscher & Malenka, 2012]. Fig. 3 shows the effects of withdrawal from oxycodone SA on the expression of GluN subunit mRNAs. There were no significant changes in the expression of *GluN1* [F_(3, 31)_ =0.9200, p = 0.4384] or *GluN2A* [F_(3, 32)_=1.111, p=0.3590] mRNA at 2-h after last oxycodone intake (Figs. 3A, C). *GluN2B* mRNA expression is significantly downregulated [F_(3, 30)_=3.255, p=0.0353] at 2-h in the LgA-H group compared to saline only (Fig. 3E). *GluN2C* mRNA levels are significantly decreased [F_(3, 28)_=2.959, p=0.0494] in the LgA-H group compared to the ShA and the control groups (Fig. 3G). *GluN2D* expression is unchanged [F_(3, 31)_=1.9535, p=0.1445] at 2-h after last oxycodone SA session (Fig. 3I). *GluN3B* mRNA expression is significantly downregulated [F_(3, 28)_=3.019, p=0.0464] at 2-h in both LgA groups compared to controls (Fig. 3K). *GluN1* mRNA expression was also not impacted [F_(3, 34)_=0.480, p=0.6984] after 31 days of withdrawal (Fig. 3B). *GluN2A* mRNA expression was significantly increased [F_(3, 32)_=4.529, p = 0.0093] in the LgA groups compared to the ShA group (Fig. 3D). *GluN2B* mRNA expression is significantly upregulated [F_(3, 32)_=4.340, p=0.0113] in LgA groups compared to the saline controls (Fig. 3F). *GluN2C* mRNA is significantly increased [F_(3, 32)_=3.307, p=0.0325] in the LgA-L group compared to the LgA-H group and the controls (Fig. 3H). *GluN2D* mRNA is significantly increased [F_(3, 33)_=2.778, p = 0.0566] in the LgA-H group compared to the ShA and control groups (Fig. 3J). There were no significant changes in the expression of *GluN3B* [F_(3, 32)_=0.032, p=0.9923] following 31 days of withdrawal (Fig. 3L). There were no significant correlations between *GluN* subunit mRNA expression and lever pressing on withdrawal day 30.

**Figure 3:**
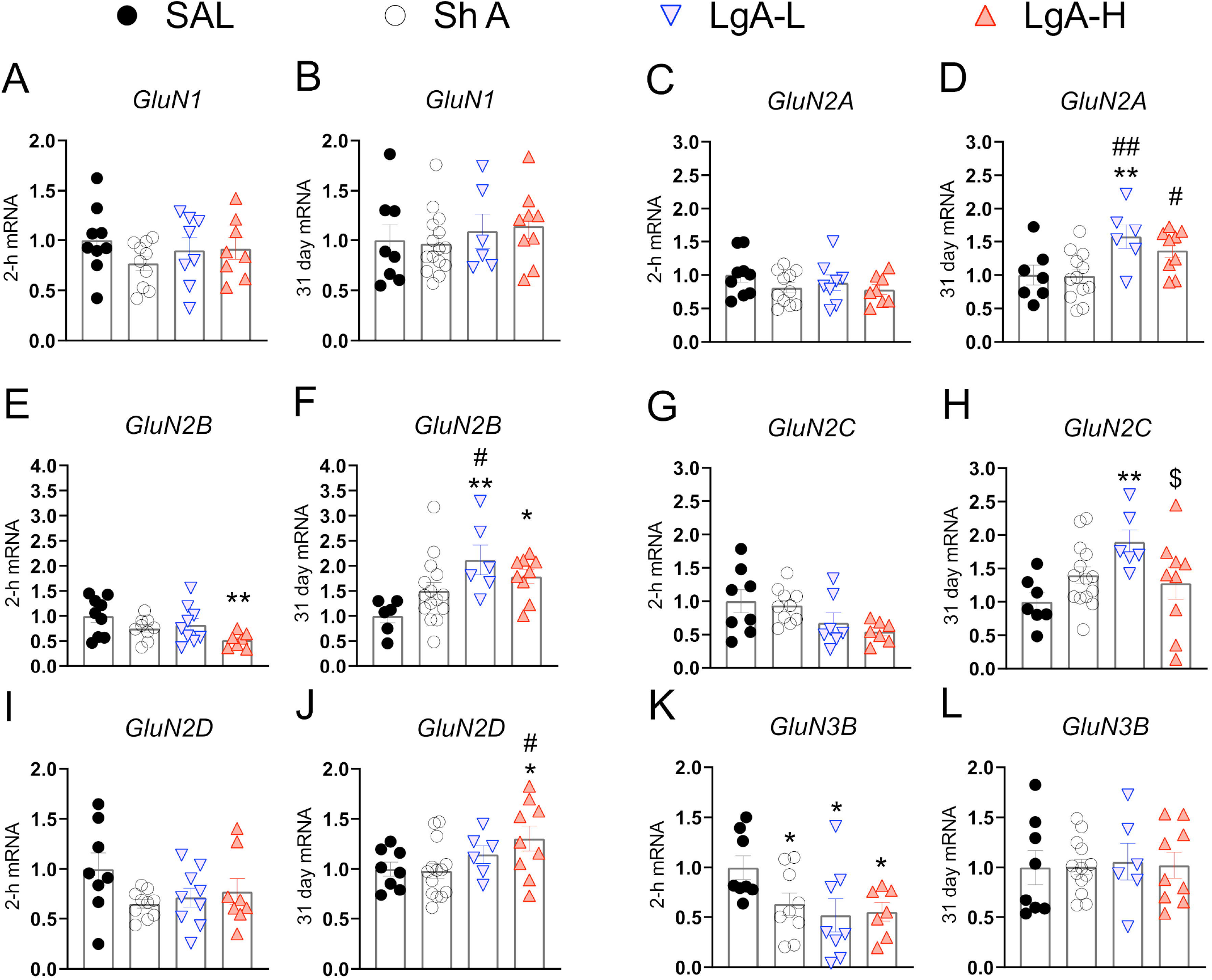
Changes in NMDA Receptor mRNA expression during oxycodone intake and withdrawal. **(A-B)** *GluN1* showed no significant changes at 2-h or 31 days. **(B-C)** *GluN2A* is significantly increased in the LgA-L group compared to the Sal and SHA groups and in the LgA-H group compared to the SHA group at 31 days. **(E)** *GluN2B* is significantly down in the LgA-H group compared to SAL at 2-h. **(F)** *GluN2B* is significantly up compared to SAL in the LgA-L and LgA-H groups and LgA-L is up compared to SHA at 31 days. **(G-H)** *GluN2C* showed no significant changes at 2-h but was significantly up in the LgA-L group compared to the SAL and LgA-H groups. **(I-J)** *GluN2D* is significantly increased in the LgA-H group compared to SAL and ShA at 31 days, but not 2-h. **(K-L)** *GluN3B* is significantly down in all groups at 2-h but not at 31 days. Key to statistics: *, **, = p < 0.05, 0.01, respectively, in comparison to Sal rats; #, ## = p < 0.05, 0.01, respectively, in comparison to ShA rats; $ = p < 0.05, respectively, in comparison to LgA-L rats. Statistical Analyses were performed as described in Fig. 2.

### 3.4 Group I Metabotropic Glutamate Receptors are Upregulated Following Abstinence from Oxycodone for 31 days

Fig. 4 shows mRNA expression data for group I metabotropic glutamate receptors. There were no significant changes in *Grm1* [F_(3, 28)_=2.526, p=0.0778] or *Grm5* [F_(3, 31)_=0.959, p=0.4245] expression at 2-h after the last oxycodone session (Figs. 4A, C). There were also no significant changes in *Grm1* after withdrawal day 31 [F_(3, 31)_=2.062, p=0.1255] (Fig. 4B). There were, however, significant increases [F_(3, 31)_=3.905, p=0.0178] in) in *Grm5* mRNA expression both LgA groups compared to the ShA group at 31 days (Fig. 4E). Interestingly, changes in *Grm1* mRNA expression were positively correlated with lever pressing on WD30 (Fig. 5C).

**Figure 4:**
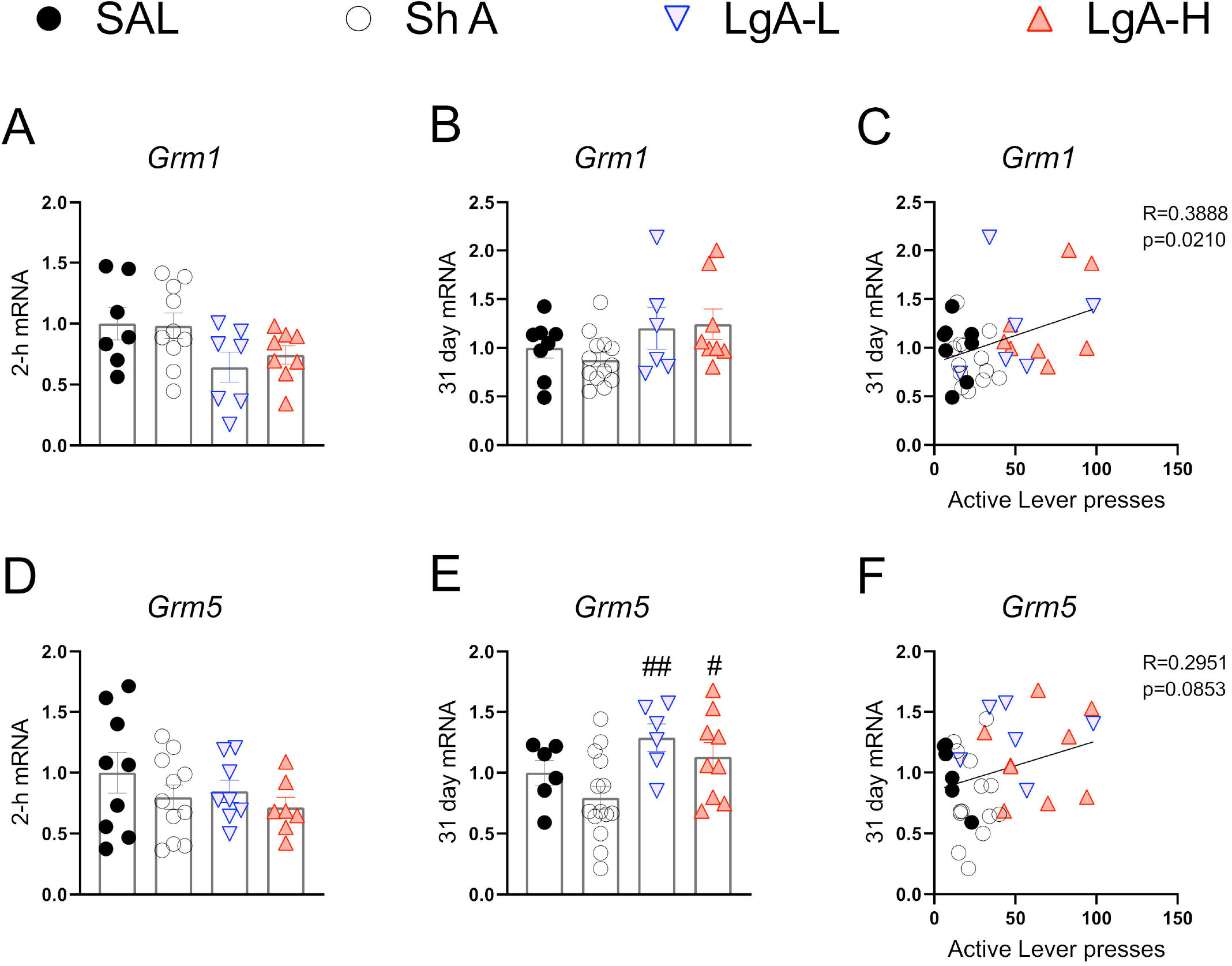
Group I metabotropic glutamate receptor mRNA expression is increased during opioid withdrawal. **(A-C)** *Grm1* showed no significant change at 2-h or 31-days after last SA, but expression at 31 days was correlated with lever pressing on WD30. **(D-F)** *Grm5* is significantly increased in both LgA groups compared to SHA group at 31 days, but mRNA expression at 31 days was not correlated with lever pressing on WD30. Key to statistics: #, ## = p < 0.05, 0.01, respectively, in comparison to ShA rats. Statistical Analyses were performed as described in Fig. 2.

### 3.5 Oxycodone SA is Associated with Downregulated Expression of Hippocampal Group II Metabotropic Glutamate Receptors

Fig. 5 shows mRNA expression data for group II metabotropic glutamate receptor mRNAs. *Grm2* [F_(3, 30)_=13.484, p < 0.0001] and *Grm3* [F_(3, 30)_=4.922, p=0.0067] mRNA levels were significantly downregulated in all oxycodone groups compared to controls at 2-h after the last oxycodone session (Figs. 5A, D). At 31 days of withdrawal, there were no significant changes in *Grm2* expression [F_(3, 31)_ = 0.06, p = 0.9618] (Fig. 5B). In contrast, *Grm3* mRNA expression remains significantly downregulated [F_(3, 31)_ = 11.991, p < 0.0001] in all oxycodone groups (Fig. 5E). Interestingly, changes in *Grm3* expression showed significant negative correlation with lever pressing on withdrawal day 31 (Fig. 5F).

**Figure 5:**
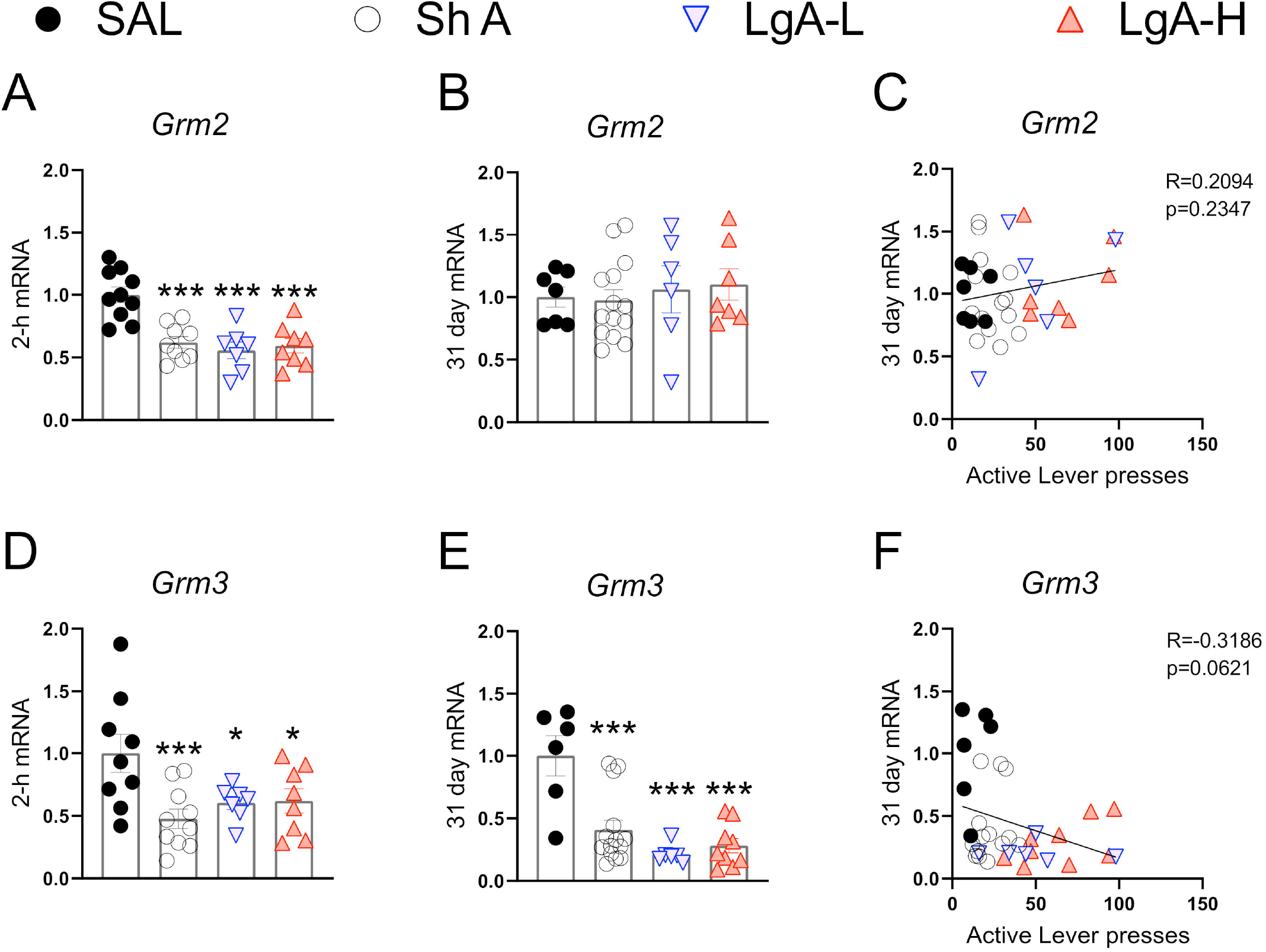
Group II metabotropic glutamate receptor mRNA expression is downregulated during oxycodone intake and maintained through long-term withdrawal. **(A-C)** *Grm2* was significantly decreased in the ShA, LgA-L, and LgA-H groups compared to SAL at 2-h but not at 31days and there was no significant correlation between lever pressing on WD30 and gene expression at 31 days. **(D-F)** Grm3 was significantly downregulated in all groups at both 2-hr and 31 days butB showed no significant correlation. Key to statistics: *, ***, = p < 0.05, 0.001, respectively, in comparison to Sal rats. Statistical Analyses were performed as described in Fig. 2.

### 3.6 Group III Metabotropic Glutamate Receptors are Upregulated in the Hippocampus Following Oxycodone Withdrawal

Fig. 6 show mRNA expression data for group III metabotropic glutamate receptors (Alexander et al., 2020). These receptors including mGluR6 receptors occur in the brain (Huang et al., 2012; Palazzo et al, 2020) and serve to suppress glutamate release [Niswender and Conn, 2010; Schoepp, 2001].There were no changes in *Grm4* [F(3, 30)=2.372, p=0.0901], *Grm6* [F(3, 32)=0.2012, p=8948], *Grm7* [F(3, 31)=2.631, p=0.0675], or *Grm8* [F(3, 32)=1.275, p=0.2997] (Figs. 6A, D, G, and J) at 2-h after the last oxycodone SA session.

**Figure 6:**
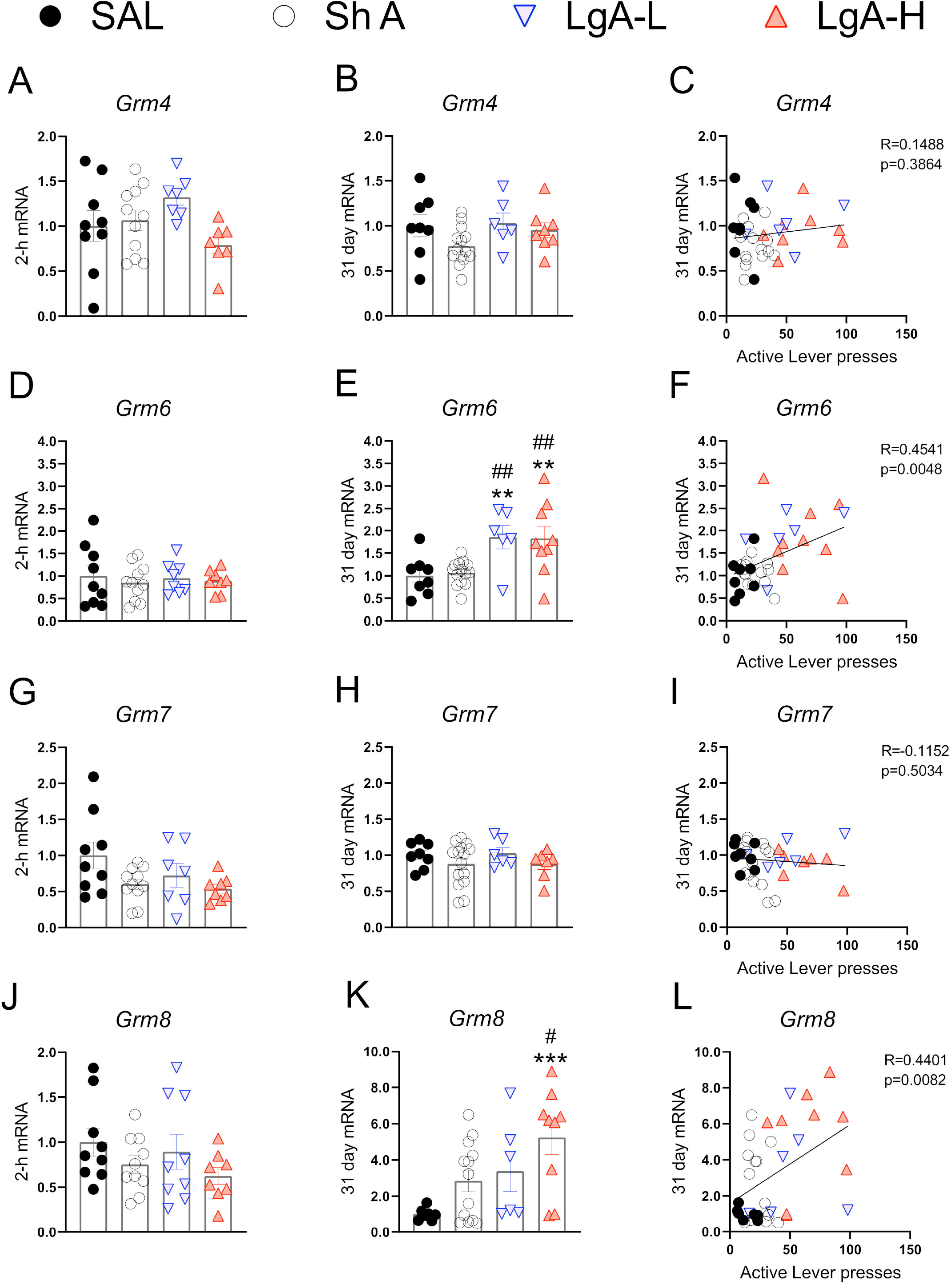
Group III metabotropic glutamate receptor mRNA expression is upregulated during withdrawal from oxycodone. **(A-C)** There were no significant changes in mRNA expression at 2-h or 31 days and no significant correlation between 31-day mRNA expression and lever pressing on WD30. (**D-F**) *Grm6* is significantly upregulated in both LgA groups compared to ShA and SAL at 31 days and there is a significant positive correlation between lever pressing on WD30 and *Grm6* expression at 31 days. **(G-I)** There were no significant changes in *Grm7*. **(J-L)** *Grm8* is significantly upregulated in the LgA-H group compared to ShA and SAL at 31 days and is positively correlated with lever pressing on WD30. Key to statistics: **, **, = p < 0.01, 0.001, respectively, in comparison to Sal rats; #, ## = p < 0.05, 0.01, respectively, in comparison to ShA rats. Statistical Analyses were performed as described in Fig. 2.

There were also no significant changes in *Grm*4 [F(3,32)=2.108, p=0.1187] during late withdrawal (Fig. 6B). However, there were significant increases in *Grm6* mRNA expression [F(3, 33)=6.634, p=0.0012] in both LgA groups compared to the other groups (Fig. 6E). *Grm7* mRNA expression shows no significant changes [F(3,33)=0.882, p=0.4608] at that time (Fig. 6H) whereas *Grm8* mRNA expression was significantly increased [F(3, 31)=4.822, p=0.0072] in the LgA-H group compared to controls (Fig. 6K). Changes in *Grm6* (Fig. 6F) and GRM8 (Fig. 6L) mRNA levels were both positively correlated with lever pressing measured on WD30.

## 4 Discussion

Oxycodone misuse and its many medical complications have made significant contributions to the current opioid public health crisis in the USA [Boscarino et al., 2010, Rudd et al., 2016]. Although efforts have been made to develop more effective treatments against opioid addiction, much more remains to be done in order to understand the biochemical and molecular effects of chronic exposure to and withdrawal from opioid drugs on the brain. Towards that end, we have been investigating the biochemical and molecular consequences of exposure to oxycodone in a rat model of drug self-administration [Blackwood et al., 2018]. We found that rats given LgA to oxycodone escalate in their drug taking and show incubation of drug seeking following a 31-day withdrawal period [Blackwood et al., 2018]. We have also shown that oxycodone SA-related behaviors are associated with significant changes in the expression of opioid receptor genes in the dorsal striatum and hippocampus of these rats [Blackwood et al., 2018]. The observation of increased oxycodone drug seeking after 31 days of withdrawal had suggested the possibility that genes involved in hippocampal memory processes might also be impacted in these animals. We thus tested the possibility that the mRNA expression of AMPAR subunits, NMDAR subunits, and metabotropic glutamate receptors might be altered in the hippocampi of rats that were exposed to oxycodone at two time points following withdrawal. We found, importantly, that most of the changes in gene expression occurred after WD30 as reported in the results.

The observations of increased expression of AMPAR subunits, *GluA1*, *GluA2*, and *GluA3* mRNA levels in the LgA-H groups that showed incubation of oxycodone craving suggest the potential involvement of these glutamate receptors in the incubation phenomenon. This suggestion is supported, in part, by the fact that changes in *GluA1*, *GluA2*, and *GluA3* mRNA levels were positively correlated with increased lever pressing (incubation) after WD30. Given the well-established role of AMPA receptors in synaptic plasticity [Diering et al, 2018], our results suggest a mechanism via which increased expression of AMPA receptors might enhance cue-induced oxycodone seeking because of strengthened synaptic connections in the hippocampus during long-term withdrawal from oxycodone. The proposition of the involvement of these receptors in oxycodone craving is also consistent with previous studies that had reported increased expression of AMPA receptors in the nucleus accumbens of rats had exhibited incubation of cocaine [Conrad et al., 2008; Mameli et al., 2009; McCutcheon et al., 2011] or methamphetamine [Murray et al 2019; Scheyer et al, 2016] craving. It is important to note that although the relative changes in expression of *GluA1*, *GluA2*, and *GluA3* were approximately the same in the LgA groups, there still might be distinct changes in receptor compositions that occur after translation and/or during assembly of these receptors that cannot be assessed by measuring only gene expression. These changes may include increased differential expression of homomeric GluA1 AMPARs (GluA2-lacking) that are calcium permeable [Hollman et al., 1991]. These compositions are known to be accompanied by enhanced AMPAR neurotransmission [Churchill et al., 1999; Mameli et al, 2009]. Importantly, Ping et al. [2008] reported that injections of antisense oligonucleotides directed against GluA1 into the NAc attenuated cocaine-primed reinstatement. Therefore, it is not far-fetched to suggest that differential compositions of AMPA receptors might modify hippocampal programs that might enhance oxycodone seeking during prolonged withdrawal from oxycodone self-administration. This discussion supports the need to examine changes in protein compositions of hippocampal AMPA receptors in follow-up studies of oxycodone SA and withdrawal. When taken together with the studies of the incubation phenomenon in rats that self-administered cocaine [Conrad et al., 2008] or methamphetamine [Scheyer et al., 2016], the present study suggests a potential role of anti-AMPAR receptor drugs in the treatment of substance use disorders to prevent relapses.

In addition to the changes in expression of *GluA* subunits in the hippocampus, we also measured potential changes in other glutamate receptors in the hippocampus after oxycodone withdrawal. It has indeed been suggested that NMDA receptor subunits might play some roles in various aspects of addiction [Hopf, 2017; Smaga et al., 2019]. We found few changes in the expression of *GluN* mRNAs in rats euthanized at the 2-hr time point. However, when compared to control and ShA rats, there were significant increases in *GluN2A* and *GluN2B* mRNA levels in both LgA groups that showed incubation of oxycodone seeking. These observations are consistent with suggestions that NMDA receptors participate in the behavioral effects of alcohol [Morisot and Ron, 2017]. Specifically, Follesa and Ticku [1995] had reported increased *GluN2A* and *GluN2B* in the hippocampi of rats chronically administered ethanol. Kalluri et al. [1998] also reported increased GluN2A and GluN2B protein levels after chronic alcohol. However, because these changes were measured only for 48 hours after cessation of alcohol intake, it is not clear what would have happened after 30 days of withdrawal. Our results are also consistent with those of Ma et al. [2007] who reported that intra-hippocampal injection of a GluN2B inhibitor, ifenprodil, was able to attenuate morphine-induced reinstatement of extinguished morphine conditioned place preference. Escalating doses of cocaine also caused increased GluN*2B* mRNA and protein levels in the hippocampus of mice engaged in a CPP paradigm [Liddie and Itzhak, 2016]. Withdrawal from cocaine self-administration is accompanied by increased GluN2A protein expression but a potential relationship between these changes and cocaine seeking was not discussed [Pomierny-Chamiolo et al., 2015]. Studies investigating the role of other GluN subunits are very scarce and the relationship of changes in NMDA receptor compositions to cue- or context-induced drug seeking remain to be fully investigated, a line of queries that might prove to be potentially fruitful.

Our study also documented some changes in the expression of metabotropic receptors during withdrawal from oxycodone self-administration. Of the type I metabotropic glutamate receptors, *Grm5* mRNA levels were increased at WD30 without there being any relationship to incubation of oxycodone seeking. Although chronic intrathecal injections of morphine also caused increased mGluR5 protein expression in the frontal cortex of mice euthanized after the last of 5 injections [Huang et al., 2019], studies on the role of this subunit on cue-induced drug seeking is non-existent. We also documented decreased mRNA levels in *Grm2* and *Grm3* mRNA levels in all rats exposed to oxycodone, suggesting profound inhibitory effects of oxycodone on those subunits. The effects of oxycodone on *Grm2* mRNA were long-lasting since they were still present even after WD30. The fact that *Grm3* mRNA levels returned to normal levels suggests that the two genes are regulated differentially by oxycodone. Our findings are consistent with the report of decreased GRM2/3 protein expression in the nucleus accumbens following withdrawal from repeated subcutaneous injection of morphine [Qian et al 2019]. Because GRM2/3 receptors are located predominantly on pre-synaptic axonal domains and glutamate terminals in the hippocampus [Ohishi et al., 1998; Petralia et al., 1996] and serve to suppress glutamate release [Niswender and Conn, 2010; Schoepp, 2001], it is possible that oxycodone-induced decrease in the expression might be compensatory in response to oxycodone-associated increased glutamate release during the drug SA experiment. Interestingly, activation of mGLu2/3 receptors by their agonist, LY379268, has been reported to attenuate reinstatement of cue-induced heroin seeking [Bossert et al., 2005]. Moreover, activation of group II metabotropic hippocampal glutamate receptors can attenuate cue-induced seeking in rats trained to self-administer ethanol [Zhao et al. 2008]. The positive modulation of GRM2/3, LY37968, also reduced cue-induced methamphetamine seeking after prolonged withdrawal (Kufahl et al, 2013). Together, these studies implicate GRM2/3 in the molecular mechanisms involved in promoting relapse after abstinence from drug taking.

We found that *Grm8* mRNA levels were increased in the LgA-H rats whereas *Grm6* was increased in all the LgA rats. Similar to group II metabotropic receptors, group III mGluRs, including GRM6 (Huang et al., 2012; Palazzo et al, 2020), are located mainly in presynaptic active zones in the brain [Ferraguti and Shigemoto, 2006; Mercier and Lodge, 2014]. The recent review [Palazzo et al., 2020] of GRM6 expression provides details about its presence beyond the visual system [Vardi et al., 2000]. The increased expression of *Grm6* mRNA levels in the LgA rats and the relationship of these increases to oxycodone seeking cement an important role for these receptors in relapse to oxycodone abuse. The increases in *Grm8* mRNA expression also correlated with incubation of oxycodone craving, thus implicating both members of metabotropic glutamate type III receptors in that behavioral phenomenon. Specific genetic manipulations of GRM6 and GRM8 should help to establish the extent to which these genes are involved in either cue- or context-induced drug seeking. Although there are, at present, very few studies have investigated potential roles of these type III metabotropic receptors in animal models of addiction, our data are consistent with those of Nielsen et al. [2008] who were able to provide evidence that metabotropic receptors, GRM6 and GRM8, were correlated with the risk of developing heroin addiction in a genome-wide association study of 110 heroin addicted individuals.

In summary, we found that there were significant changes in the expression of several glutamate receptors in the hippocampus and that some of these changes correlated positively with increased oxycodone seeking within their same individual cages at WD30, a phenomenon that may reflect relapse potential in humans under similar conditions. Because the hippocampus plays an important role in the induction of context-associated drug seeking in animal models of psychostimulants and opioids [Bossert et al., 2016; Felipe et al, 2021; Galinato et al., 2018; Noe et al., 2019; Taubenfeld et al., 2010], it will be important to investigate context- and cue-induced in parallel to assess if similar or distinct molecular changes are associated with these behavioral phenomena. In addition, although we have discussed the molecular changes in terms of their facilitating oxycodone drug seeking behaviors, it is possible that these changes might have actually been consequences to lever pressing. We think that this is unlikely because we euthanized the rats 24 hours after the last drug seeking test. Moreover, our results are consistent with those of other investigators who have implicated some of these glutamate receptors in mediating drug seeking behaviors [Wolf, 2016]. Nevertheless, it will be important to investigate the effects of prolonged drug withdrawal in the absence of drug seeking tests. In any case, our results are consistent with the proposal that glutamatergic and memory systems might play important roles in the manifestations and clinical course of opioid use disorders [Heinsbroek et al., 2020]. The present observations broadened our insight into potential ways that glutamate receptors might act to promote incubation of oxycodone seeking after prolonged withdrawal. Dissecting these mechanisms better should help the development of novel targets for oxycodone addiction. When taken together with previous results with cocaine and methamphetamine withdrawal, our observations hint to the use of AMPAR antagonist and mGluR agonist in a general approach to therapeutic interventions against substance use disorders (SUDs).

## Author contributions

C.A.B performed self-administration experiments. A.J.S. Performed RT-PCR experiments. A.J.S., C.A.B., J.L.C prepared manuscript. J.L.C supervised the overall project.

## Acknowledgments

The authors are grateful to several reviewers whose suggestions helped us to write a better manuscript.

## References

Alexander SPH, Christopoulos A, Davenport AP, Kelly E, Mathie A, Peters JA, Veale EL, Armstrong JF, Faccenda E, Harding SD, Pawson AJ, Sharman JL, Southan C, Davies JA; CGTP Collaborators. THE CONCISE GUIDE TO PHARMACOLOGY 2019/20: G protein-coupled receptors. Br J Pharmacol. 2019 Dec;176 Suppl 1(Suppl 1):S21–S141. doi: 10.1111/bph.14748. PMID: 31710717; PMCID: PMC6844580.

Allegri N, Mennuni S, Rulli E, Vanacore N, Corli O, Floriani I, De Simone I, Allegri M, Govoni S, Vecchi T, Sandrini G, Liccione D, Biagioli E. Systematic Review and Meta-Analysis on Neuropsychological Effects of Long-Term Use of Opioids in Patients With Chronic Noncancer Pain. Pain Pract. 2019 Mar;19(3):328–343. doi: 10.1111/papr.12741. Epub 2018 Dec 10. PMID: 30354006.

Blackwood CA, Hoerle R, Leary M, Schroeder J, Job MO, McCoy MT, Ladenheim B, Jayanthi S, Cadet JL. Molecular Adaptations in the Rat Dorsal Striatum and Hippocampus Following Abstinence-Induced Incubation of Drug Seeking After Escalated Oxycodone Self-Administration. Mol Neurobiol. 2019 May;56(5):3603–3615. doi: 10.1007/s12035-018-1318-z. Epub 2018 Aug 28. PMID: 30155791; PMCID: PMC6477015.

Blackwood, C.A., Leary, M., Salisbury, A., McCoy, M.T., and Cadet, J.L. (2019b). Escalated Oxycodone Self-Administration Causes Differential Striatal mRNA Expression of FGFs and IEGs Following Abstinence-Associated Incubation of Oxycodone Craving. Neuroscience 415, 173–183. doi: https://doi.org/10.1016/j.neuroscience.2019.07.030.

Blackwood CA, McCoy MT, Ladenheim B, Cadet JL. Escalated Oxycodone Self-Administration and Punishment: Differential Expression of Opioid Receptors and Immediate Early Genes in the Rat Dorsal Striatum and Prefrontal Cortex. Front Neurosci. 2020 Jan 9;13:1392. doi: 10.3389/fnins.2019.01392. PMID: 31998063; PMCID: PMC6962106.

Boscarino JA, Rukstalis M, Hoffman SN, Han JJ, Erlich PM, Gerhard GS, Stewart WF. Risk factors for drug dependence among out-patients on opioid therapy in a large US health-care system. Addiction. 2010 Oct;105(10):1776–82. doi: 10.1111/j.1360-0443.2010.03052.x. Epub 2010 Aug 16. PMID: 20712819.

Bossert JM, Adhikary S, St Laurent R, Marchant NJ, Wang HL, Morales M, Shaham Y. Role of projections from ventral subiculum to nucleus accumbens shell in context-induced reinstatement of heroin seeking in rats. Psychopharmacology (Berl). 2016 May;233(10):1991–2004. doi: 10.1007/s00213-015-4060-5. Epub 2015 Sep 7. PMID: 26344108; PMCID: PMC4781679.

Bossert JM, Busch RF, Gray SM. The novel mGluR2/3 agonist LY379268 attenuates cue-induced reinstatement of heroin seeking. Neuroreport. 2005 Jun 21;16(9):1013–6. doi: 10.1097/00001756-200506210-00026. PMID: 15931079.

Boudreau AC, Wolf ME. Behavioral sensitization to cocaine is associated with increased AMPA receptor surface expression in the nucleus accumbens. J Neurosci. 2005 Oct 5;25(40):9144–51. doi: 10.1523/JNEUROSCI.2252-05.2005. PMID: 16207873; PMCID: PMC6725751.

Brady, A.M., Saul, R.D., and Wiest, M.K. (2010). Selective deficits in spatial working memory in the neonatal ventral hippocampal lesion rat model of schizophrenia. Neuropharmacology 59(7-8), 605–611. doi: 10.1016/j.neuropharm.2010.08.012.

Cadet, J.L., Bisagno, V., and Milroy, C.M. (2014). Neuropathology of substance use disorders. Acta Neuropathologica 127(1), 91–107. doi: 10.1007/s00401-013-1221-7.

Cadet JL, Brannock C, Krasnova IN, Jayanthi S, Ladenheim B, McCoy MT, Walther D, Godino A, Pirooznia M, Lee RS. Genome-wide DNA hydroxymethylation identifies potassium channels in the nucleus accumbens as discriminators of methamphetamine addiction and abstinence. Mol Psychiatry. 2017 Aug;22(8):1196–1204. doi: 10.1038/mp.2016.48. Epub 2016 Apr 5. PMID: 27046646; PMCID: PMC7405865.

Cadet JL, Brannock C, Ladenheim B, McCoy MT, Krasnova IN, Lehrmann E, Becker KG, Jayanthi S. Enhanced upregulation of CRH mRNA expression in the nucleus accumbens of male rats after a second injection of methamphetamine given thirty days later. PLoS One. 2014 Jan 27;9(1):e84665. doi: 10.1371/journal.pone.0084665. PMID: 24475032; PMCID: PMC3903495.

Chambers, R.A., and Taylor, J.R. (2004). Animal modeling dual diagnosis schizophrenia: Sensitization to cocaine in rats with neonatal ventral hippocampal lesions. Biological Psychiatry 56(5), 308–316. doi: https://doi.org/10.1016/j.biopsych.2004.05.019.

Chen ZG, Liu X, Wang W, Geng F, Gao J, Gan CL, Chai JR, He L, Hu G, Zhou H, Liu JG. Dissociative role for dorsal hippocampus in mediating heroin self-administration and relapse through CDK5 and RhoB signaling revealed by proteomic analysis. Addict Biol. 2017 Nov;22(6):1731–1742. doi: 10.1111/adb.12435. Epub 2016 Aug 22. PMID: 27549397.

Churchill L, Swanson CJ, Urbina M, Kalivas PW. Repeated cocaine alters glutamate receptor subunit levels in the nucleus accumbens and ventral tegmental area of rats that develop behavioral sensitization. J Neurochem. 1999 Jun;72(6):2397–403. doi: 10.1046/j.1471-4159.1999.0722397.x. PMID: 10349849.

Conrad KL, Tseng KY, Uejima JL, Reimers JM, Heng LJ, Shaham Y, Marinelli M, Wolf ME. Formation of accumbens GluR2-lacking AMPA receptors mediates incubation of cocaine craving. Nature. 2008 Jul 3;454(7200):118–21. doi: 10.1038/nature06995. Epub 2008 May 25. PMID: 18500330; PMCID: PMC2574981.

Daiwile AP, Jayanthi S, Ladenheim B, McCoy MT, Brannock C, Schroeder J, Cadet JL. Sex Differences in Escalated Methamphetamine Self-Administration and Altered Gene Expression Associated With Incubation of Methamphetamine Seeking. Int J Neuropsychopharmacol. 2019 Nov 1;22(11):710–723. doi: 10.1093/ijnp/pyz050. PMID: 31562746; PMCID: PMC6902093.

Diering, G.H., and Huganir, R.L. (2018). The AMPA Receptor Code of Synaptic Plasticity. Neuron 100(2), 314–329. doi: https://doi.org/10.1016/j.neuron.2018.10.018.

Elahi-Mahani, A., Heysieattalab, S., Hosseinmardi, N., Janahmadi, M., Seyedaghamiri, F., and Khoshbouei, H. (2018). Glial cells modulate hippocampal synaptic plasticity in morphine dependent rats. Brain Research Bulletin 140, 97–106. doi: https://doi.org/10.1016/j.brainresbull.2018.04.006.

Felipe JM, Palombo P, Bianchi PC, Zaniboni CR, Anésio A, Yokoyama TS, Engi SA, Carneiro-de-Oliveira PE, Planeta CDS, Leão RM, Cruz FC. Dorsal hippocampus plays a causal role in context-induced reinstatement of alcohol-seeking in rats. Behav Brain Res. 2021 Feb 1;398:112978. doi: 10.1016/j.bbr.2020.112978. Epub 2020 Oct 24. PMID: 33169700.

Ferraguti F, Shigemoto R. Metabotropic glutamate receptors. Cell Tissue Res. 2006 Nov;326(2):483–504. doi: 10.1007/s00441-006-0266-5. Epub 2006 Jul 18. PMID: 16847639.

Follesa P, Ticku MK. Chronic ethanol treatment differentially regulates NMDA receptor subunit mRNA expression in rat brain. Brain Res Mol Brain Res. 1995 Mar;29(1):99–106. doi: 10.1016/0169-328x(94)00235-7. PMID: 7770006.

Fuchs RA, Evans KA, Ledford CC, Parker MP, Case JM, Mehta RH, See RE. The role of the dorsomedial prefrontal cortex, basolateral amygdala, and dorsal hippocampus in contextual reinstatement of cocaine seeking in rats. Neuropsychopharmacology. 2005 Feb;30(2):296–309. doi: 10.1038/sj.npp.1300579. PMID: 15483559.

Galinato MH, Takashima Y, Fannon MJ, Quach LW, Morales Silva RJ, Mysore KK, Terranova MJ, Dutta RR, Ostrom RW, Somkuwar SS, Mandyam CD. Neurogenesis during Abstinence Is Necessary for Context-Driven Methamphetamine-Related Memory. J Neurosci. 2018 Feb 21;38(8):2029–2042. doi: 10.1523/JNEUROSCI.2011-17.2018. Epub 2018 Jan 23. Erratum in: J Neurosci. 2019 Oct 2;39(40):7992. PMID: 29363584; PMCID: PMC5824740.

Glick, S.D., and Cox, R.D. (1978). Changes in morphine self-administration after tel-diencephalic lesions in rats. Psychopharmacology 57(3), 283–288. doi: 10.1007/BF00426752.

Heinsbroek JA, De Vries TJ, Peters J. Glutamatergic Systems and Memory Mechanisms Underlying Opioid Addiction. Cold Spring Harb Perspect Med. 2020 May 27:a039602. doi: 10.1101/cshperspect.a039602. Epub ahead of print. Erratum in: Cold Spring Harb Perspect Med. 2020 Jun 1;10(6): PMID: 32341068; PMCID: PMC7718856.

Hopf FW. Do specific NMDA receptor subunits act as gateways for addictive behaviors? Genes Brain Behav. 2017 Jan;16(1):118–138. doi: 10.1111/gbb.12348. Epub 2016 Nov 18. PMID: 27706932; PMCID: PMC5351810.

Hollmann M, Hartley M, Heinemann S. Ca2+ permeability of KA-AMPA--gated glutamate receptor channels depends on subunit composition. Science. 1991 May 10;252(5007):851–3. doi: 10.1126/science.1709304. PMID: 1709304.

Huang M, Luo L, Zhang Y, Wang W, Dong J, Du W, Jiang W, Xu T. Metabotropic glutamate receptor 5 signalling induced NMDA receptor subunits alterations during the development of morphine-induced antinociceptive tolerance in mouse cortex. Biomed Pharmacother. 2019 Feb;110:717–726. doi: 10.1016/j.biopha.2018.12.042. Epub 2018 Dec 13. PMID: 30553198.

Huang, Y. Y., Haug, M. F., Gesemann, M., & Neuhauss, S. C. (2012). Novel expression patterns of metabotropic glutamate receptor 6 in the zebrafish nervous system. PLoS One, 7(4), e35256. doi:10.1371/journal.pone.0035256

Jury, N. J., Radke, A. K., Pati, D., Kocharian, A., Mishina, M., Kash, T. L., & Holmes, A. (2018). NMDA receptor GluN2A subunit deletion protects against dependence-like ethanol drinking. Behav Brain Res, 353, 124–128. doi:10.1016/j.bbr.2018.06.029

Kalluri HS, Mehta AK, Ticku MK. Up-regulation of NMDA receptor subunits in rat brain following chronic ethanol treatment. Brain Res Mol Brain Res. 1998 Jul 15;58(1-2):221–4. doi: 10.1016/s0169-328x(98)00112-0. PMID: 9685652.

Koob, G.F., and Volkow, N.D. (2010). Neurocircuitry of Addiction. Neuropsychopharmacology 35(1), 217–238. doi: 10.1038/npp.2009.110.

Kroll, S.L., Nikolic, E., Bieri, F., Soyka, M., Baumgartner, M.R., and Quednow, B.B. (2018). Cognitive and socio-cognitive functioning of chronic non-medical prescription opioid users. Psychopharmacology (Berl) 235(12), 3451–3464. doi: 10.1007/s00213-018-5060-z.

Kufahl PR, Watterson LR, Nemirovsky NE, Hood LE, Villa A, Halstengard C, Zautra N, Olive MF. Attenuation of methamphetamine seeking by the mGluR2/3 agonist LY379268 in rats with histories of restricted and escalated self-administration. Neuropharmacology. 2013 Mar;66:290–301. doi: 10.1016/j.neuropharm.2012.05.037. Epub 2012 May 31. PMID: 22659409; PMCID: PMC3442155.

LeGates TA, Kvarta MD, Tooley JR, Francis TC, Lobo MK, Creed MC, Thompson SM. Reward behaviour is regulated by the strength of hippocampus-nucleus accumbens synapses. Nature. 2018 Dec;564(7735):258–262. doi: 10.1038/s41586-018-0740-8. Epub 2018 Nov 26. PMID: 30478293; PMCID: PMC6292781.

Liddie S, Itzhak Y. Variations in the stimulus salience of cocaine reward influences drug-associated contextual memory. Addict Biol. 2016 Mar;21(2):242–54. doi: 10.1111/adb.12191. Epub 2014 Oct 28. PMID: 25351485; PMCID: PMC4412748.

Lüscher, C., and Malenka, R.C. (2012). NMDA receptor-dependent long-term potentiation and long-term depression (LTP/LTD). Cold Spring Harbor perspectives in biology 4(6), a005710. doi: 10.1101/cshperspect.a005710.

Ma YY, Chu NN, Guo CY, Han JS, Cui CL. NR2B-containing NMDA receptor is required for morphine-but not stress-induced reinstatement. Exp Neurol. 2007 Feb;203(2):309–19. doi: 10.1016/j.expneurol.2006.08.014. Epub 2006 Oct 2. PMID: 17014848.

Mameli M, Halbout B, Creton C, Engblom D, Parkitna JR, Spanagel R, Lüscher C. Cocaine-evoked synaptic plasticity: persistence in the VTA triggers adaptations in the NAc. Nat Neurosci. 2009 Aug;12(8):1036–41. doi: 10.1038/nn.2367. Epub 2009 Jul 13. PMID: 19597494.

McCutcheon JE, Wang X, Tseng KY, Wolf ME, Marinelli M (2011). Calcium-permeable AMPA receptors are present in nucleus accumbens synapses after prolonged withdrawal from cocaine self-administration but not experimenter-administered cocaine. J Neurosci. 2011 Apr 13;31(15):5737–43. doi: 10.1523/JNEUROSCI.0350-11.2011. PMID: 21490215; PMCID: PMC3157976.

Mercier MS, Lodge D. Group III metabotropic glutamate receptors: pharmacology, physiology and therapeutic potential. Neurochem Res. 2014 Oct;39(10):1876–94. doi: 10.1007/s11064-014-1415-y. Epub 2014 Aug 22. PMID: 25146900.

Monyer H, Sprengel R, Schoepfer R, Herb A, Higuchi M, Lomeli H, Burnashev N, Sakmann B, Seeburg PH. Heteromeric NMDA receptors: molecular and functional distinction of subtypes. Science. 1992 May 22;256(5060):1217–21. doi: 10.1126/science.256.5060.1217. PMID: 1350383.

Murray CH, Loweth JA, Milovanovic M, Stefanik MT, Caccamise AJ, Dolubizno H, Funke JR, Foster Olive M, Wolf ME. AMPA receptor and metabotropic glutamate receptor 1 adaptations in the nucleus accumbens core during incubation of methamphetamine craving. Neuropsychopharmacology. 2019 Aug;44(9):1534–1541. doi: 10.1038/s41386-019-0425-5. Epub 2019 May 30. PMID: 31146278; PMCID: PMC6785134.

Nielsen DA, Ji F, Yuferov V, Ho A, Chen A, Levran O, Ott J, Kreek MJ. Genotype patterns that contribute to increased risk for or protection from developing heroin addiction. Mol Psychiatry. 2008 Apr;13(4):417–28. doi: 10.1038/sj.mp.4002147. Epub 2008 Jan 15. PMID: 18195715; PMCID: PMC3810149.

Niswender CM, Conn PJ. Metabotropic glutamate receptors: physiology, pharmacology, and disease. Annu Rev Pharmacol Toxicol. 2010;50:295–322. doi: 10.1146/annurev.pharmtox.011008.145533. PMID: 20055706; PMCID: PMC2904507.

Noe E, Bonneau N, Fournier ML, Caillé S, Cador M, Le Moine C. Arc reactivity in accumbens nucleus, amygdala and hippocampus differentiates cue over context responses during reactivation of opiate withdrawal memory. Neurobiol Learn Mem. 2019 Mar;159:24–35. doi: 10.1016/j.nlm.2019.02.007. Epub 2019 Feb 13. PMID: 30771462.

Nomura, A., Shigemoto, R., Nakamura, Y., Okamoto, N., Mizuno, N., & Nakanishi, S. (1994). Developmentally regulated postsynaptic localization of a metabotropic glutamate receptor in rat rod bipolar cells. Cell, 77(3), 361–369. doi:https://doi.org/10.1016/0092-8674(94)90151-1

Ohishi H, Neki A, Mizuno N. Distribution of a metabotropic glutamate receptor, mGluR2, in the central nervous system of the rat and mouse: an immunohistochemical study with a monoclonal antibody. Neurosci Res. 1998 Jan;30(1):65–82. doi: 10.1016/s0168-0102(97)00120-x. PMID: 9572581.

Olive, M.F. (2009). Metabotropic glutamate receptor ligands as potential therapeutics for addiction. Curr Drug Abuse Rev 2(1), 83–98. doi: 10.2174/1874473710902010083.

Ortiz J, Harris HW, Guitart X, Terwilliger RZ, Haycock JW, Nestler EJ. Extracellular signal-regulated protein kinases (ERKs) and ERK kinase (MEK) in brain: regional distribution and regulation by chronic morphine. J Neurosci. 1995 Feb;15(2):1285–97. doi: 10.1523/JNEUROSCI.15-02-01285.1995. PMID: 7532701; PMCID: PMC6577831.

Palazzo E, Boccella S, Marabese I, Pierretti G, Guida F, Maione S. The Cold Case of Metabotropic Glutamate Receptor 6: Unjust Detention in the Retina? Curr Neuropharmacol. 2020;18(2):120–125. doi: 10.2174/1570159X17666191001141849. PMID: 31573889; PMCID: PMC7324884.

Petralia RS, Wang YX, Niedzielski AS, Wenthold RJ. The metabotropic glutamate receptors, mGluR2 and mGluR3, show unique postsynaptic, presynaptic and glial localizations. Neuroscience. 1996 Apr;71(4):949–76. doi: 10.1016/0306-4522(95)00533-1. PMID: 8684625.

Ping A, Xi J, Prasad BM, Wang MH, Kruzich PJ. Contributions of nucleus accumbens core and shell GluR1 containing AMPA receptors in AMPA- and cocaine-primed reinstatement of cocaine-seeking behavior. Brain Res. 2008 Jun 18;1215:173–82. doi: 10.1016/j.brainres.2008.03.088. Epub 2008 Apr 13. PMID: 18486119; PMCID: PMC2728035.

Pomierny-Chamiolo L, Miszkiel J, Frankowska M, Pomierny B, Niedzielska E, Smaga I, Fumagalli F, Filip M. Withdrawal from cocaine self-administration and yoked cocaine delivery dysregulates glutamatergic mGlu5 and NMDA receptors in the rat brain. Neurotox Res. 2015 Apr;27(3):246–58. doi: 10.1007/s12640-014-9502-z. Epub 2014 Nov 19. PMID: 25408547; PMCID: PMC4353866.

Portugal GS, Al-Hasani R, Fakira AK, Gonzalez-Romero JL, Melyan Z, McCall JG, Bruchas MR, Morón JA. Hippocampal long-term potentiation is disrupted during expression and extinction but is restored after reinstatement of morphine place preference. J Neurosci. 2014 Jan 8;34(2):527–38. doi: 10.1523/JNEUROSCI.2838-13.2014. PMID: 24403152; PMCID: PMC3870935.

Qian Z, Wu X, Qiao Y, Shi M, Liu Z, Ren W, Han J, Zheng Q. Downregulation of mGluR2/3 receptors during morphine withdrawal in rats impairs mGluR2/3- and NMDA receptor-dependent long-term depression in the nucleus accumbens. Neurosci Lett. 2019 Jan 18;690:76–82. doi: 10.1016/j.neulet.2018.10.018. Epub 2018 Oct 11. PMID: 30315852.

Riley, J., Eisenberg, E., Müller-Schwefe, G., Drewes, A.M., and Arendt-Nielsen, L. (2008). Oxycodone: a review of its use in the management of pain. Current Medical Research and Opinion 24(1), 175–192. doi: 10.1185/030079908X253708.

Rogers, J.L., and See, R.E. (2007). Selective inactivation of the ventral hippocampus attenuates cue-induced and cocaine-primed reinstatement of drug-seeking in rats. Neurobiol Learn Mem 87(4), 688–692. doi: 10.1016/j.nlm.2007.01.003.

Rudd, R.A., Aleshire, N., Zibbell, J.E., and Matthew Gladden, R. (2016). Increases in Drug and Opioid Overdose Deaths—United States, 2000–2014. American Journal of Transplantation 16(4), 1323–1327. doi: 10.1111/ajt.13776.

Scheyer AF, Loweth JA, Christian DT, Uejima J, Rabei R, Le T, Dolubizno H, Stefanik MT, Murray CH, Sakas C, Wolf ME. AMPA Receptor Plasticity in Accumbens Core Contributes to Incubation of Methamphetamine Craving. Biol Psychiatry. 2016 Nov 1;80(9):661–670. doi: 10.1016/j.biopsych.2016.04.003. Epub 2016 Apr 12. PMID: 27264310; PMCID: PMC5050076.

Schoepp DD. Unveiling the functions of presynaptic metabotropic glutamate receptors in the central nervous system. J Pharmacol Exp Ther. 2001 Oct;299(1):12–20. PMID: 11561058.

Smaga I, Sanak M, Filip M. Cocaine-induced Changes in the Expression of NMDA Receptor Subunits. Curr Neuropharmacol. 2019;17(11):1039–1055. doi: 10.2174/1570159X17666190617101726. PMID: 31204625; PMCID: PMC7052821.

Substance-Related and Addictive Disorders in Diagnostic and Statistical Manual of Mental Disorders.).

Taubenfeld SM, Muravieva EV, Garcia-Osta A, Alberini CM. Disrupting the memory of places induced by drugs of abuse weakens motivational withdrawal in a context-dependent manner. Proc Natl Acad Sci U S A. 2010 Jul 6;107(27):12345–50. doi: 10.1073/pnas.1003152107. Epub 2010 Jun 21. PMID: 20566855; PMCID: PMC2901477.

Terman, G.W., Wagner, J.J., and Chavkin, C. (1994). Kappa opioids inhibit induction of long-term potentiation in the dentate gyrus of the guinea pig hippocampus. The Journal of Neuroscience 14(8), 4740. doi: 10.1523/JNEUROSCI.14-08-04740.1994.

Vardi N, Duvoisin R, Wu G, Sterling P. Localization of mGluR6 to dendrites of ON bipolar cells in primate retina. J Comp Neurol. 2000 Jul 31;423(3):402–12. doi: 10.1002/1096-9861(20000731)423:3<402::aid-cne4>3.0.co;2-e. PMID: 10870081.

Wang JQ, Tang Q, Parelkar NK, Liu Z, Samdani S, Choe ES, Yang L, Mao L. Glutamate signaling to Ras-MAPK in striatal neurons: mechanisms for inducible gene expression and plasticity. Mol Neurobiol. 2004 Feb;29(1):1–14. doi: 10.1385/MN:29:1:01. PMID: 15034219.

Wang N, Ge F, Cui C, Li Y, Sun X, Sun L, Wang X, Liu S, Zhang H, Liu Y, Jia M, Yang M. Role of Glutamatergic Projections from the Ventral CA1 to Infralimbic Cortex in Context-Induced Reinstatement of Heroin Seeking. Neuropsychopharmacology. 2018 May;43(6):1373–1384. doi: 10.1038/npp.2017.279. Epub 2017 Nov 14. PMID: 29134962; PMCID: PMC5916356.

Wilson N, Kariisa M, Seth P, Smith H IV, Davis NL. (2020) Drug and Opioid-Involved Overdose Deaths — United States, 2017–2018. MMWR Morb Mortal Wkly Rep 69:290–297. DOI: http://dx.doi.org/10.15585/mmwr.mm6911a4external icon

Wolf ME. Synaptic mechanisms underlying persistent cocaine craving. Nat Rev Neurosci. 2016 Jun;17(6):351–65. doi: 10.1038/nrn.2016.39. Epub 2016 May 6. PMID: 27150400; PMCID: PMC5466704.

Zhang, Y., Loh, H.H., and Law, P.Y. (2016). Effect of Opioid on Adult Hippocampal Neurogenesis. ScientificWorldJournal 2016, 2601264. doi: 10.1155/2016/2601264.

Zhao, Y., Dayas, C. V., Aujla, H., Baptista, M. A. S., Martin-Fardon, R., & Weiss, F. (2006). Activation of group II metabotropic glutamate receptors attenuates both stress and cue-induced ethanol-seeking and modulates c-fos expression in the hippocampus and amygdala. The Journal of neuroscience: the official journal of the Society for Neuroscience, 26(39), 9967–9974. doi:10.1523/JNEUROSCI.2384-06.2006

Zoubková, H., Tomášková, A., Nohejlová, K., Černá, M., & Šlamberová, R. (2019). Prenatal Exposure to Methamphetamine: Up-Regulation of Brain Receptor Genes. Frontiers in neuroscience, 13, 771–771. doi:10.3389/fnins.2019.00771

